# Early-life PFAS exposure induces systemic metabolic alterations in plasma and brain in mice

**DOI:** 10.64898/2026.10.14.758497

**Authors:** Tuulia Hyötyläinen, Daniel Duberg, Ilia Evstafev, Alex M. Dickens, Eriika Savontaus, Matej Orešič

## Abstract

Per- and polyfluoroalkyl substances (PFAS) are persistent environmental contaminants associated with adverse health effects, including emerging neurodevelopmental and neurological outcomes. Although PFAS can cross the placental and blood–brain barriers, the metabolic mechanisms underlying their neurotoxicity and potential sex-specific effects remain poorly understood. Here, we investigated the effects of prenatal and early-life exposure to a human-relevant PFAS mixture on plasma and brain metabolism in mice. The mixture contained perfluorooctanesulfonic acid (PFOS), perfluorooctanoic acid (PFOA), perfluorohexanesulfonic acid (PFHxS), perfluorodecanoic acid (PFDA), and perfluorononanoic acid (PFNA) at environmentally relevant, subtoxic concentrations.

Mass spectrometry–based metabolomics revealed pronounced, sex-dependent metabolic alterations in both plasma and brain. Female brains exhibited more extensive metabolic changes, particularly in lipid and arachidonic acid metabolism, steroid hormone pathways, and carbohydrate and energy metabolism. In contrast, male brains showed more selective perturbations, predominantly involving amino acid and neuroactive signaling pathways related to neurotransmitter metabolism. These distinct metabolic signatures indicate that early-life PFAS exposure affects brain metabolism through sex-dependent mechanisms. Our findings identify the brain metabolome as a sensitive target of early-life PFAS exposure and emphasize the importance of considering sex as a biological variable when assessing the neurotoxicity of persistent environmental contaminants.

## 1. Introduction

Per- and polyfluoroalkyl substances (PFAS) are a diverse class comprising thousands of synthetic fluorinated compounds that have been used for decades in industrial applications and consumer products due to their oil- and water-repellent properties. Their high chemical stability, driven by the strength of the carbon–fluorine bond, renders them extremely resistant to environmental and biological degradation. This persistence, combined with extensive global production and historically limited regulatory oversight, has led to widespread environmental contamination and near-universal human exposure. Drinking water, food, dust, and consumer products represent dominant exposure pathways.

The toxicological and epidemiological evidence linking PFAS to adverse health effects has expanded substantially. In adults, numerous cohort and biomonitoring studies have associated PFAS exposure with cancer, dyslipidaemia, immune suppression, liver and kidney dysfunction, reproductive impairments, and developmental toxicity.^1–13^ More recently, concerns about neurological and neurodevelopmental consequences have intensified as new data emerge from both human cohorts and experimental models.^1, 14^ ^15^ Observational studies increasingly report associations between perinatal or childhood PFAS exposure and attention-deficit/hyperactivity disorder (ADHD), autism spectrum disorder (ASD), reduced cognitive performance, behavioral dysregulation, and altered motor development, conditions that represent a substantial global public health burden.^16, 17^ We have recently shown that prenatal PFAS exposure predicts multimodal brain structural and functional outcomes in children aged 5 years.^18^

Mechanistically, growing evidence suggests that PFAS exposure during critical windows of development may directly influence central nervous system (CNS) maturation.^19^ PFAS readily cross the placental barrier, and emerging evidence indicates that several emerging or replacement PFAS exhibit even higher transplacental transfer efficiencies than legacy chemicals such as PFOS and PFOA, with their branched chain isomers showing particularly efficient placental passage.^20–22^ Following entry into fetal circulation, PFAS can cross the blood–brain barrier (BBB) and accumulate in the brain, with high concentrations being found in the brain stem, hippocampus, hypothalamus, pons/medulla and thalamus.^1^ This type of regional accumulation of PFAS within the brain suggests that certain brain structures may be disproportionately affected. Although the BBB becomes more selective with maturation, it remains relatively permeable during early development, increasing susceptibility to chemical penetration.

The fetal and early postnatal periods represent highly sensitive windows for neurodevelopment, during which neural proliferation, migration, synaptogenesis, myelination, and metabolic programming are occurring at rapid rates ^23^.^24, 25^ Disruption of these processes, whether through endocrine activity, altered lipid metabolism, mitochondrial dysfunction, neuroinflammation, interference with neurotransmitter systems, or dysregulation of gut-liver-brain axis - can have lasting consequences for brain function.^14, 15, 26, 27^ Indeed, developmental exposure to PFAS has been linked to long-lasting neurotoxic effects, including impaired neuronal differentiation, disrupted synaptic plasticity, and cognitive deficits .^28, 29^

PFAS have been implicated to disturb several adverse pathways, including modulation of peroxisome proliferator-activated receptors (PPARs), interference with calcium signaling, disturbance of thyroid hormone homeostasis, induction of oxidative stress, and alterations in glucose and lipid metabolic pathways, all of which may influence brain development and metabolism.^14, 27, 30, 31^ Specific effects include altered glutamate and tryptophan metabolism, changes in calcium homeostasis, and dysregulation of neuroactive pathways like cholinergic and GABAergic synapses, leading to neurotoxicity. The neurotoxicity of PFAS has been suggested to involve both direct effects on brain structure and function and indirect effects on other organs that affect brain homeostasis.^14^ Exposure to PFAS has been observed to activate major pathogenic mechanisms linked with oxidative stress, mitochondrial dysfunction, mitochondrial bioenergetics, and DNA damage.^32^

Together, this growing body of evidence indicates that PFAS may adversely affect brain metabolism and neurodevelopment, however, the exact mechanisms are poorly understood. Particularly, there is very limited information of potential metabolic changes in the brain due to PFAS exposure. A recent study in male mice showed that a mixture of PFAS caused suppression of purine metabolism and activation of arachidonic acid metabolism, suggesting involvement in neurotoxic effects.^15^ Another study showed that PFOA exposure neurotransmitters were altered, as well as metabolites related to lipid metabolism.^33^ In a more targeted study, metabolite changes were observed in cortex, mainly perturbation of cortical lipids, such as phosphatidylcholines and sphingomyelins, as well as multiple amino acids.^34^ There is a need for experimental studies that can clarify how these chemicals alter metabolic programmes in the developing brain. Understanding these pathways is essential not only for risk assessment but also for informing regulatory decisions and identifying vulnerable populations.

In this study, we investigated the impact of prenatal and early-life exposure of PFAS on plasma and brain metabolic profiles in mice, with a particular focus on brain-related alteration and sex-specific responses. The PFAS mixture consisted of perfluorooctanesulfonic acid (PFOS), PFOA, perfluorohexanesulfonic acid (PFHxS), perfluorodecanoic acid (PFDA), and perfluorononanoic acid (PFNA), with concentrations and concentration ratios mimicking human-relevant exposures. We performed comprehensive metabolomic characterization of murine plasma and brain samples by applying three complementary analytical platforms. The primary aim was to evaluate subtoxic, environmentally relevant exposure levels that reflect real-world conditions.

## 2. Methods

### 2.1. Chemicals

All solvents were HPLC grade or LC-MS grade, from Honeywell (Morris Plains, NJ, USA), Fisher Scientific (Waltham, MA, USA) or Sigma-Aldrich (St. Louis, MO, USA). Mass spectrometry grade ammonium acetate and reagent grade formic acid were also from Sigma-Aldrich (St. Louis, MO, USA). The lipid standards were from Avanti Polar Lipids Inc. (Alabaster, AL, USA).. One native performance standard, 7H-dodecafluoroheptanoic acid, was purchased from ABCR (Karlsruhe, Germany). Cholic acid (CA), chenodeoxycholic acid (CDCA), deoxycholic acid (DCA), dehydrocholic Acid (DHCA), glycocjolic acid (GCA), glyco chenodeoxycholic acid (GCDCA), lithocholic acid (LCA), taurocholic acid (TCA), taurochenodeoxycholic acid (TCDCA), taurodeoxycholic acid TDCA, taurodehydrocholic acid (TDHCA), taurohyocholic acid (THCA), taurohyodeoxycholic acid (THDCA), taurolithocholic acid (TLCA), and tauroursocholic acid (TUDCA) were obtained from Sigma-Aldrich (St. Luis, MO, USA), hyodeoxycholic acid (HDCA), hyocholic acid (HCA), murocholic acids α, β and ω (αMCA, βMCA, ωMCA), 7- oxohyodeoxycholic acid (7-oxo-HDCA), 7-oxodeoxycholic acid (7-oxo-DCA), 12-oxo lithocholic acid (12-oxo-LCA), tauromurocholic acid), murocholic acids α, β and ω (TαMCA, TβMCA, TωMCA), glycodehydrocholic Acid (GDHCA), glycohyocholic acid (GHCA), and glycohyodeoxycholic acid) GHDCA from Steraloids (Newport, RI, U.S.A), glycolithocholic acid (GLCA) and glycoursocholic acid (GUDCA) from Calbiochem (Gibbstown, NJ, U.S.A), and glycodeoxycholic acid (GDCA) and ursocholic acid (UDCA) from Fluka (Buchs, Switzerland). Internal standards CA-d4, LCA-d4, UDCA-d4, CDCA-d4, DCA-d4, GCA-d4, GLCA-d4, GUDCA-d4 and GCDCA-d4 were obtained from Qmx laboratories Ltd. (Essex, UK). For quality assurance (QA), standard reference material serum NIST SRM 1950 (for lipidomics and metabolomics) and 1957 (for PFAS and bile acids) was purchased from the National Institute of Standards and Technology (NIST) at the US Department of Commerce (Washington, DC, USA).

### 2.2. Murine exposure study

The experimental procedures were approved by the Animal Experiment Board of Regional State Administrative Agency for Southern Finland (license ESAVI/45421/2022). Animal care was conducted in accordance with the Directive 2010/63/EU of the European Parliament on the protection of animals used for scientific purposes and with the institutional and national guidelines for the care and use of laboratory animals. NOD.scid mice were purchased from Charles River UK limited, Margata, England, and the strains were maintained in Central Animal Laboratory of University of Turku. Mice were group-housed on a fixed 12-hour light/dark cycle and at a constant temperature of 21±3°C, with free access to irradiated chow (Teklad 2916, Envigo Bioproducts Inc) and UV-treated tap water unless otherwise indicated.

Female NOD mice were randomized into control and exposure groups. The PFAS exposure *via* drinking water was started 4-6 days after conformation of pregnancy and continued through gestation and early life period until 8 weeks of age. At 8 weeks of age, 8 offspring were sacrificed in the control group (3 female mice and 6 male mice) along with 8 offspring from the exposure group (4 female mice and 6 male mice).

Plasma and brain samples were collected and stored at -80 °C until chemical analyses. Samples were prepared and analyzed in randomized order. There was no difference is size of brains between the control and exposed mice.

For preparation of vehicle and dosing solution two concentrated solutions were prepared.

**50% sucrose solution in tap water** was prepared by dissolving 400g of D(+)-Sucrose (Saccharose) (VWR Cat. N 27480.294) in 800 mL water.

**Solution of PFAS mixture** was prepared by dissolving a mixture of PFAS in 2 L of water resulting in PFAS solution as shown in Table 1.

**Table 1.** PFAS solution used in the exposure.

| Compound | mass, mg | μmols |
| --- | --- | --- |
| PFOS | 78.2 | 156.36 |
| PFOA | 46.2 | 111.58 |
| PFHxS | 9.6 | 23.99 |
| PFNA | 8.2 | 17.67 |
| PFDA | 4.8 | 9.34 |

**Table 2.** LC-MS conditions for the two methods used in the study.

| Conditions | Polar/semipolar compounds:<br>Method 1 | Lipidomics: Method 2 | HILIC: Method 3 |
| --- | --- | --- | --- |
| **Injection volume** | 2 µl | 1 µl | 2 µl |
| **Column** | C18 precolumn (Waters Corporation, Wexford, Ireland) and an inline filter, pore size 0,2 µm (Waters Corporation, Wexford, Ireland). + ACQUITY UPLC® BEH C18 column (2.1 mm × 100 mm, particle size 1.7 µm) by Waters (Milford, MA, USA) | C18 precolumn (Waters Corporation, Wexford, Ireland) and an inline filter, pore size 0,2 µm (Waters Corporation, Wexford, Ireland).+ ACQUITY UPLC® BEH C18 column (2.1 mm × 100 mm, particle size 1.7 µm) by Waters (Milford, MA, USA) | XBridge™ Premier BEH Amide 2.5 µm (2.1 mm x 100 mm) |
| **Mobile phases** | A H<sub>2</sub>O:MeOH (v/v 70:30) with 2 mM ammonium acetate | A 10 mM ammonium acetate and 0.1% Formic Acid in H<sub>2</sub>O | Mobile phase A: 10 mM NH<sub>4</sub>HCO<sub>2</sub> in H<sub>2</sub>O and 0.125% formic acid |
| | B MeOH with containing 2 mM ammonium acetate | B Acetonitrile:Isopropanol (v/v 1:1) with 0.1% Formic Acid and 10 mM ammonium acetate | Mobile phase B: 10 mM $\text{NH}_4\text{HCO}_2$ in ACN:H<sub>2</sub>O (95:5) and 0.125% formic acid |
| **Gradient** | • 0-1.5 min: B was increased from 5% to 30%<br>• 1.5-4.5 min,<br>• B increased to 70%;<br>• 4.5-7.5 min,<br>• B increased to 100% and held for 5.5 min.<br>• A post-time of 6 minutes | • 0-2 min, B was increased from 35% to 80%<br>• 2-7 min, B increased to 100%<br>• 7-14 min, B was held at 100%.<br>• A post-time of 7 min | • 0-7.7 min: 0% A 100% B<br>• 7.7-9.5 min: 30% A 70% B<br>• 9.50-10.25 min: 60% A 40% B<br>• 10.25-12.75 min: 70% A 30% B<br>• 12.75-17.00 min: 0% A 100% B |
| **Flow rate** | 0.4 mLmin<sup>-1</sup> | 0.4 mLmin<sup>-1</sup> | 0.4 mLmin<sup>-1</sup> |
| **MS conditions** | Dual ESI ionization source with capillary voltage 4.5 kV, nozzle voltage 1500 V, N<sub>2</sub> pressure in the nebulized was 21 psi and the N<sub>2</sub> flow rate and temperature as sheath gas was 11 Lmin<sup>-1</sup> and 379 °C, respectively. Drying gas flow was set to 10 Lmin<sup>-1</sup> and | Dual ESI ionization source with capillary voltage 3.64 kV, nozzle voltage 1500 V, N<sub>2</sub> pressure in the nebulized was 21 psi and the N<sub>2</sub> flow rate and temperature as sheath gas was 11 Lmin<sup>-1</sup> and 379 °C, respectively. Drying gas flow was set to 10 Lmin<sup>-1</sup> and temperature to 193 °C. | Dual ESI ionization source with capillary voltage 3.64 kV, nozzle voltage 1500 V, N<sub>2</sub> pressure in the nebulized was 21 psi and the N<sub>2</sub> flow rate and temperature as sheath gas was 11 Lmin<sup>-1</sup> and 379 °C, respectively. Drying gas flow was set to 10 Lmin<sup>-1</sup> and |
|  | temperature to 150 °C. m/z range 100-1700 in negative ion mode. | m/z range 100-1700 in positive ion mode | temperature to 193 °C. m/z range 50-1200 in negative ion mode. |

Vehicle preparation was done by adding 1mL of the 50% sucrose solution to 100 mL of water and mixed. PFAS administration solution was done by mixing 1 mL of 50% sucrose solution, 5 mL of PFAS mixture solution and 100 mL of water.

### 2.3. LC-MS analysis

All samples were randomized before sample preparation and analysis. The samples were analyzed using three parallel methods, one aimed at analysis of semipolar metabolites, the second one for the analysis of lipids and the third one for polar metabolites.

All analyses were done by an ultra-high-performance liquid chromatography quadrupole time-of-flight mass spectrometry (UHPLC-QTOFMS). For methods 1 and 3, , the UHPLC coupled with an Xevo G3 QTOF system (Waters Corporation, Milford, Massachusetts, USA). The data acquisition was done using MassLynx V.4.2 (Waters Corporation). For Method 2 the UHPLC system used in this work was a 1290 Infinity II system from Agilent Technologies (Santa Clara, CA, USA). The system was equipped with a multi sampler (maintained at 10 °C), a quaternary solvent manager and a column thermostat (maintained at 50 °C). MassHunter B.06.01 software (Agilent Technologies, Santa Clara, CA, USA) was used for all data acquisition

#### 2.3.1. Sample preparation

Brain samples were weighted, and 0.9% NaCl was added so that the ratio of tissue to buffer was 1 mg tissue to 5 µL buffer and the samples were homogenized manually. 40 µl of the homogenate was extracted with 480 µL of monophasic MeOH:MTBE:IPA (20:15:15, v/v) extraction solvent. After vortex-mixing (30 s), the extract was kept on an ice plate (45 min) and then transferred to a tube fitted with a 0.45 µm filter and centrifuged at 3000 × g for 3 min. For exposomics/metabolomics analysis, a 100 µL aliquot was transferred to a vial, evaporated in a SpeedVac system, and reconstituted with 50 µL MeOH:H2O (7:3, v/v) for analysis.

#### 2.3.2. Analysis of PFAS, polar and semipolar metabolites (Methods 1 and 3)

Quantitation was done using 6-point calibration (bile acids c= 20-640 ng/mL, polar metabolites c=0.1 -80 μg/mL, PFAS c= 3.75-120 ng/mL). Quantification of other bile acids was done using the following compounds: Chenodeoxycholic acid (CDCA), Cholic acid (CA), Deoxycholic acid (DCA), Glycochenodeoxycholic acid (GCDCA), Glycocholic acid (GCA), Glycodehydrocholic acid (GDCA), Glycodeoxycholic acid (GDCA), Glycohyocholic acid (GHCA), Glycohyodeoxycholic acid (GHDCA), Glycolithocholic acid (GLCA), Glycoursodeoxycholic acid (GUDCA), Hyocholic acid (HCA), Hyodeoxycholic acid (HDCA), Litocholic acid (LCA), alpha-Muricholic acid (αMCA), Tauro-alpha- muricholic acid (T-α-MCA), Tauro-beta-muricholic acid(T-β-MCA), Taurochenodeoxycholic acid (TCDCA), Taurocholic acid (TCA), Taurodehydrocholic acid (THCA), Taurodeoxycholic acid (TDCA), Taurohyodeoxycholic acid (THDCA), Taurolithocholic acid (TLCA), Tauro-omega-muricholic acid (TωMCA) and Tauroursodeoxycholic acid (TDCA) and polar metabolites was done using alanine, citric acid, fumaric acid, glutamic acid, glycine, lactic acid, malic acid, 2-hydroxybutyric acid, 3-hydroxybutyric acid, linoleic acid, oleic acid, palmitic acid, stearic acid, cholesterol, fructose, glutamine, indole-3-propionic acid, isoleucine, leucine, proline, succinic acid, valine, asparagine, aspartic acid, arachidonic acid, glycerol-3-phosphate, lysine, methionine, ornithine, phenylalanine, serine and threonine. PFAS standards included PFOS, PFOA, PFHxS, PFDA and PFNA. The curves had R values >0.98 for most of the compounds. Identification was done based on in-house library (m/z, MS/MS, retention times) that is based on analysis of authentic standards.

Standard solutions extracted blanks (n=3), pooled QC samples (n=3, an aliquot of each sample pooled), in-house serum QC and NIST SRM 1950 (human plasma) were analyzed together with the samples. Identification was done based on in-house library with retention time and spectral data (level 1 identification, based on Metabolic Standard Initiative^35^). For pooled samples, identified metabolites had an average RSD of 21.1%.

#### 2.3.3. Analysis of molecular lipids (Method 2)

Calibration curves using 1-hexadecyl-2-(9Z-octadecenoyl)-sn-glycero-3- phosphocholine (PC(16:0e/18:1(9Z))), 1-(1Z-octadecenyl)-2-(9Z-octadecenoyl)-sn-glycero-3-phosphocholine (PC(18:0p/18:1(9Z))), 1-stearoyl-2-hydroxy-sn-glycero-3- phosphocholine (LPC(18:0)), 1-oleoyl-2-hydroxy-sn-glycero-3-phosphocholine (LPC(18:1)), 1-palmitoyl-2-oleoyl-sn-glycero-3-phosphoethanolamine (PE(16:0/18:1)), 1- (1Z-octadecenyl)-2-docosahexaenoyl-sn-glycero-3-phosphocholine (PC(18:0p/22:6)) and 1-stearoyl-2-linoleoyl-sn-glycerol (DG(18:0/18:2)), 1-(9Z-octadecenoyl)-sn-glycero- 3-phosphoethanolamine (LPE(18:1)), N-(9Z-octadecenoyl)-sphinganine (Cer(d18:0/18:1(9Z))), 1-hexadecyl-2-(9Z-octadecenoyl)-sn-glycero-3- phosphoethanolamine (PE(16:0/18:1)) from Avanti Polar Lipids, 1-Palmitoyl-2-Hydroxy-sn-Glycero-3-Phosphatidylcholine (LPC(16:0)), 1,2,3 trihexadecanoalglycerol (TG(16:0/16:0/16:0)), 1,2,3-trioctadecanoylglycerol (TG(18:0/18:0/18:0) and 3β-hydroxy- 5-cholestene-3-stearate (ChoE(18:0)), 3β-Hydroxy-5-cholestene-3-linoleate (ChoE(18:2)) from Larodan, were prepared to the following concentration levels: 100, 500, 1000, 1500, 2000 and 5000 ng/mL (in CHCl3:MeOH, 2:1, v/v) including 1250 ng/mL of each internal standard. The curves had R values >0.99 for all lipids. Identification was done based on in-house library with retention time and spectral data (level 1 and 2 identification, based on Metabolic Standard Initiative). Lipid abbreviations are as follows: lipid headgroup followed by the fatty acyl composition (number of carbons in the fatty acyl, number of double bonds). Lipid headgroups: Ceramide (Cer), Sphingomyelin (SM), Phosphatidylcholine (PC), Alkylphosphatidylcholine (PC-O), Phosphatidylcholine plasmalogen (PC-P), Lysophosphatidylcholine (LPC), Phosphatidylethanolamine (PE), Akylphosphatidylethanolamine (PE-O), Phosphatidylethanolamine plasmalogen (PE-P), Lysophosphatidylethanolamine (LPE), Phosphatidylinositol (PI), Phosphatidylserine (PS), Cholesterol Ester (Ce), Diacylglycerol (DG), Triacylglycerol (TG).

Standard solutions extracted blanks (n=3), pooled QC samples (n=3, an aliquot of each sample pooled), in-house serum QC and NIST CRM 1950 (human plasma) were analyzed together with the samples. For pooled samples, identified lipids had an average RSD in the pooled samples of 13.9%.

### 2.4. Data preprocessing

Processing of MS data was performed using the open-source software package MZmine 4.8 ^36^. The following steps were applied in this processing: (i) Crop filtering with a m/z range of 350 – 1200 m/z and an RT range of 2.0 to 12 minutes, (ii) Mass detection with a noise level of 750, (iii) Chromatogram builder with a minimum time span of 0.08 min, minimum height of 1000 and a m/z tolerance of 0.006 m/z or 10.0 ppm, (iv) Chromatogram deconvolution using the local minimum search algorithm with a 70% chromatographic threshold, 0.05 min minimum RT range, 5% minimum relative height, 1200 minimum absolute height, a minimum ration of peak top/edge of 1.2 and a peak duration range of 0.0– - 5.0, (v), Isotopic peak grouper with a m/z tolerance of 5.0 ppm, RT tolerance of 0.05 min, maximum charge of 2 and with the most intense isotope set as the representative isotope, (vi) Peak filter with minimum 8 data points, a FWHM between 0.0 and 0.2, tailing factor between 0.45 and 2.22 and asymmetry factor between 0.40 and 2.50, (vii) Join aligner with a m/z tolerance of 0.009 or 10.0 ppm and a weight for of 2, a RT tolerance of 0.15 min and a weight of 1 and with no requirement of charge state or ID and no comparison of isotope pattern, (viii) Peak list row filter with a minimum of 10% of the samples (ix) Gap filling using the same RT and m/z range gap filler algorithm with an m/z tolerance of 0.009 m/z or 11.0 ppm, (x) Identification of lipids and metabolites using a custom database search with an m/z tolerance of 0.009 m/z or 10.0 ppm and a RT tolerance of 0.15 min, and (xi) Normalization using internal standards: For lipids: PE(17:0/17:0), SM(d18:1/17:0), Cer(d18:1/17:0), LPC(17:0), TG(17:0/17:0/17:0) and PC(16:0/d30/18:1)) for identified lipids and closest internal standard for the unknown lipids followed by calculation of the concentrations based on lipid-class concentration curves. For polar metabolites the following internal standards were used: Valine-d8, Glutamic acid-d5, Succinic acid-d4, Heptadecanoic acid, Lactic acid-d3, Citric acid-d4. 3-Hydroxybutyric acid-d4, Arginine-d7, Tryptophan-d5, Glutamine-d5, 1-D4-CA,1-D4- CDCA,1-D4-CDCA,1-D4-GCA,1-D4-GCDCA,1-D4-GLCA,1-D4-GUDCA,1-D4-LCA,1-D4-TCA, 1-D4-UDCA and closest internal standard for the unknown metabolites followed by calculation of the concentrations-based concentration curves. For data filtering, we have removed compounds that were present at blank samples (peak area >5 times that of blank) and compounds that had RSD>30% in the pooled quality control samples. MS/MS data was done for the pooled quality control samples using auto MS/MS mode. The identification was based on in-house spectral libraries, based on MS/MS and retention time data on authentic standards as well as public spectral libraries.

### 2.5. Data analysis

Data analyses were conducted using Metaboanalyst 6.0^37^ and the R statistical programming language (version 4.1.2) (https://www.r-project.org/). In order to correct for heteroscedasticity, the exposure datasets were pre-processed by log2 transformation and scaling to zero mean and unit variance (autoscaled)as metabolomics data is not normally distributed. The statistical analyses included principal component analysis (PCA), analysis of variance (ANOVA), t-test and fold-change between controls and PFAS concentrations measured, Spearman correlations, and partial correlations between the PFAS concentrations measured and metabolites.

Pathway enrichment analysis was performed using the MetaboAnalyst 6.0 web platform with the Functional Analysis (MS Peaks) module (Pang et al. 2022^38^). This approach supports functional analysis of untargeted metabolomics data generated from high-resolution mass spectrometry. The pathway analysis was done with the data of the polar and semipolar metabolites. The input data for the pathway analysis comprised complete LC-HRMS data, i.e. both identified and unknown metabolites, obtained in negative ionization mode. First, we performed statistical analyses using adjusted linear model between control and each treatment, resulting in fold change, p values and FDR values. The whole input peak list, with peak names given as their numeric mass (m/z) values for putative annotation, and the statistical results with p-values and t-score were used for the pathway analysis. Two algorithms were applied separately, namely, Mummichog and Gene Set Enrichment Analysis (GSEA) and two pathway libraries were used in the pathway analysis, namely human scale metabolic model MNF (from MetaboAnalyst Mummichog package) and Kyoto Encyclopedia of Genes and Genomes (KEGG) pathways for *Homo Sapiens* to determine the relative significance of the identified pathways (Li et al. 2020). The mass tolerance for the pathway analysis was set at 7 ppm, and we also used an advanced option to select representative adducts by removing isotopic adducts as these have been already removed in our data preprocessing step.

## 3. Results

### 3.1. Plasma and brain metabolome and PFAS

The study included 16 non-obese diabetic (NOD) mice at 8 weeks of age, 8 in non-exposed (control) group (3 female, 6 male) and eight in the PFAS-exposed group (4 female, 6 male). We identified >800 metabolites in paired murine plasma and brain samples, spanning lipids, nucleosides free fatty acids, amino acids, biogenic amines, carbohydrates, bile acids, steroids hormones, vitamins and other metabolite classes (**Figures 1** and **2**, **Supplementary Tables 1** and **2**). Among lipids, triacylglycerols (TGs), phosphatidylcholines (PCs), phosphatidylethanolamines (PEs) as well as ceramides (CERs) and sphingomyelins (SMs). were the most abundant lipid classes in the brain, whereas PCs were most abundant in plasma, followed by TGs. Only a small number of brain metabolites differed between female and male mice (**Supplementary Table 3**).

**Figure 1.**
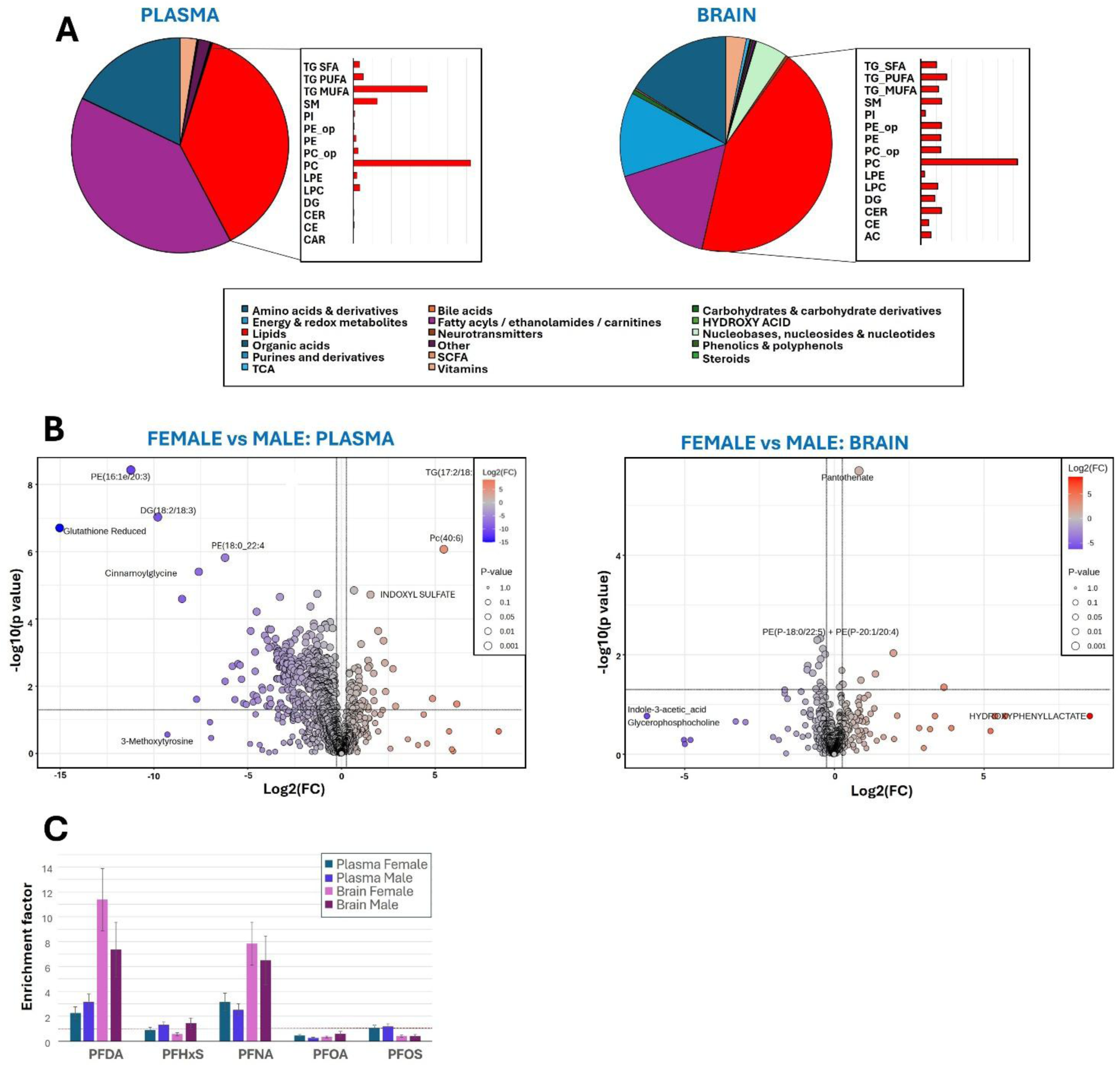
Overview of plasma and brain metabolome. (**A**) Metabolite classes present in murine plasma and brain samples from the control group. (**B**) Volcano plots of metabolic differences between female and male mice. (**C**) PFAS enrichment factor, in relation to exposure dose ratio in plasma and brain. Lipid class abbreviations: TG_SFA, TGs containing saturated fatty acids (FAs); TG_PUFA, TGs containing polyunsaturated FAs; TG_MUFA, TGs containing monounsaturated FAs; PI, phosphatidylinositol; PE_O, ether PE; PC_O/P, ether PC; LPE, lyso PE; LPC, lyso PC; DG, diacylglycerol; CE, cholesterol ester; AC, acylcarnitine.

PFAS exposure did not associate with the brain size. Overall, male mice had slightly larger brains than females (FC = 1.05, p = 0.0063). PFAS concentrations in plasma and brain are shown in **Table 3**. Female mice exhibited generally higher PFAS levels than males, both in circulation and in brain tissue. Specifically, PFDA and PFNA were higher in plasma, while PFDA, PFNA, and PFOS were higher in the brains of female mice.

**Table 3.** A) PFAS levels in the brains of female and male mice.

|  | Plasma (ng/ml) |  |  |  | Brain (ng/g) |  |  |  |
| --- | --- | --- | --- | --- | --- | --- | --- | --- |
|  | FC (F/M) | p | Female | Male | FC (F/M) | p | Female | Male |
| PFDA | 1.38 | **0.0438** | 805.5 | 858.0 | 2.23 | **0.0071** | 149.85 | 67.16 |
| PFHxS | 0.90 | 0.8476 | 1291.0 | 1432.0 | 0.57 | 0.0749 | 30.12 | 52.55 |
| PFNA | 1.64 | **0.0058** | 3830.7 | 2329.0 | 1.74 | **0.0028** | 353.32 | 202.65 |
| PFOA | 0.95 | 0.3888 | 4667.0 | 4897.8 | 0.83 | 0.3499 | 88.51 | 106.17 |
| PFOS | 1.19 | 0.1414 | 8578.2 | 7199.5 | 1.42 | **0.0261** | 172.26 | 121.56 |

To assess relative accumulation of the individual PFAS, we compared the proportional contribution of each compound in plasma and brain with its proportion in the exposure mixture. This analysis showed that PFDA and PFNA were enriched in brain tissue relative to their proportion in the exposure mixture, indicating stronger brain accumulation compared with the other PFAS. In contrast, PFOA and PFOS showed lower proportional representation in the brain relative to their proportions in the exposure mixture (**Figure 1C**).

### 3.2. PFAS exposure is associated with sex-specific changes in the metabolome

In plasma, PFAS-treated mice exhibited significant metabolic alterations (Figure 2). Principal component analysis (PCA) revealed a clear sex-specific separation between groups (Figure 2A). In male mice, PFAS treatment resulted in pronounced metabolic changes (Figure 2B, Supplementary Table X), with 389 metabolites showing significant differences (p < 0.05, FDR < 0.1), including widespread downregulation of lipids. In females, fewer metabolites showed significant changes (15 metabolites); however, this may reflect the smaller number of female mice from which plasma samples were available. In female mice, several lipids were also downregulated, while 3-methoxytyrosine was significantly upregulated (FC = 10.2). In addition to the identified metabolites and lipids, a large number of unidentified compounds showed significant differences in both female and male mouse brains.

**Figure 2.**
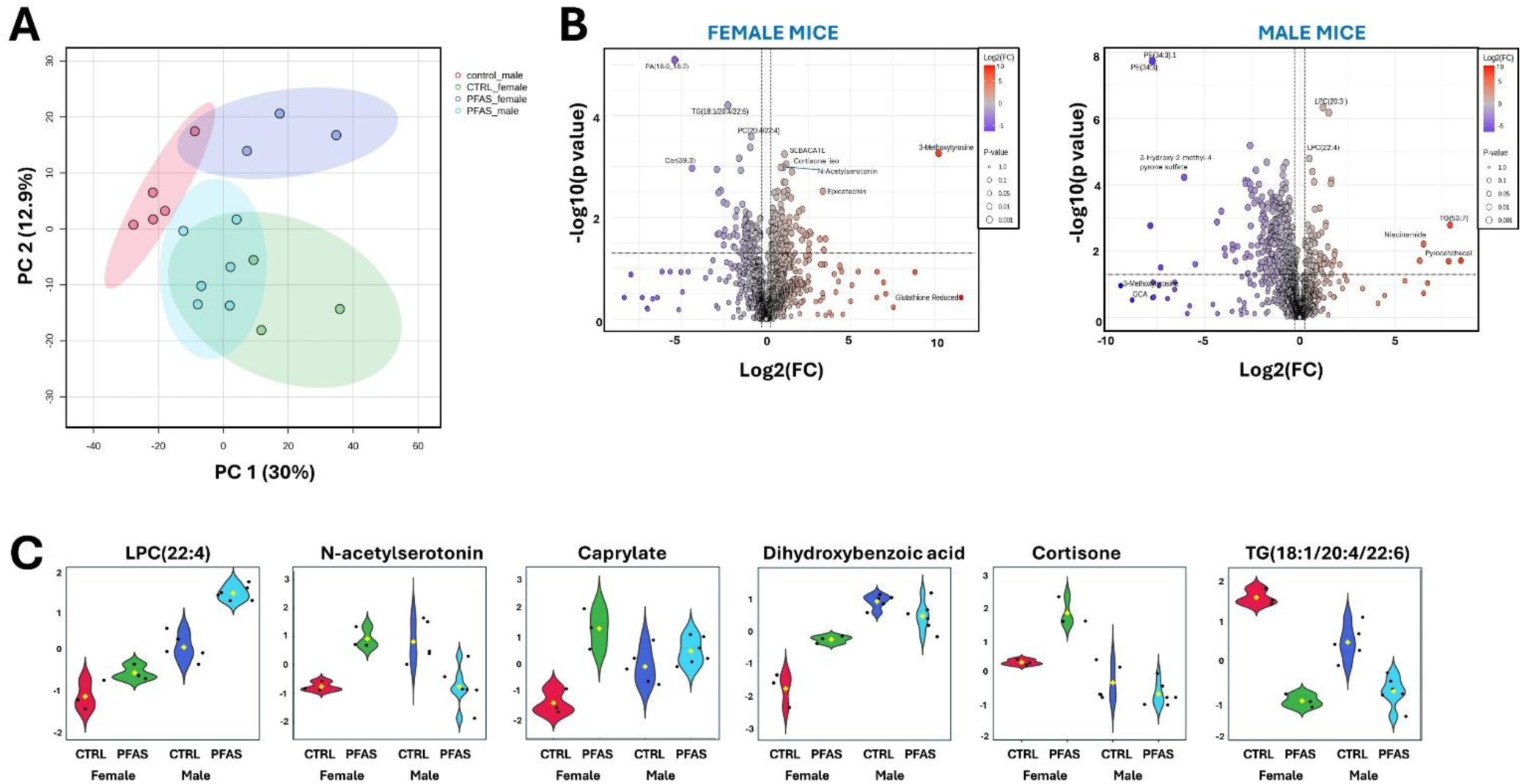
(A) Principal component analysis (PCA) of identified plasma metabolites across treatment groups and sex. **(B)** Volcano plots showing differential metabolites in female and male mice (PFAS-exposed vs control). **(C)** Selected examples of significantly altered metabolites.

In brain, PFAS exposure induced clear sex-specific effects in the mouse brain metabolome, with a markedly stronger impact observed in females. A heatmap showing fold changes between PFAS-treated mice and control mice is presented in Figures 3-4, with detailed values provided in Supplementary Table 3-4. In female mice, PFAS exposure led to upregulation of several ceramide species, lysophosphatidylcholines (LPCs), phosphatidylcholines (PCs), and phosphatidylethanolamines (PEs). In addition, free fatty acids and amino acids were increased, whereas several acylcarnitines and nucleotide derivatives were downregulated. Notably, arachidonic acid (AA) and docosahexaenoic acid (DHA) showed significant upregulation in female mouse brains, while the neuronal marker N-acetyl-L-aspartic acid was significantly downregulated.

**Figure 3.**
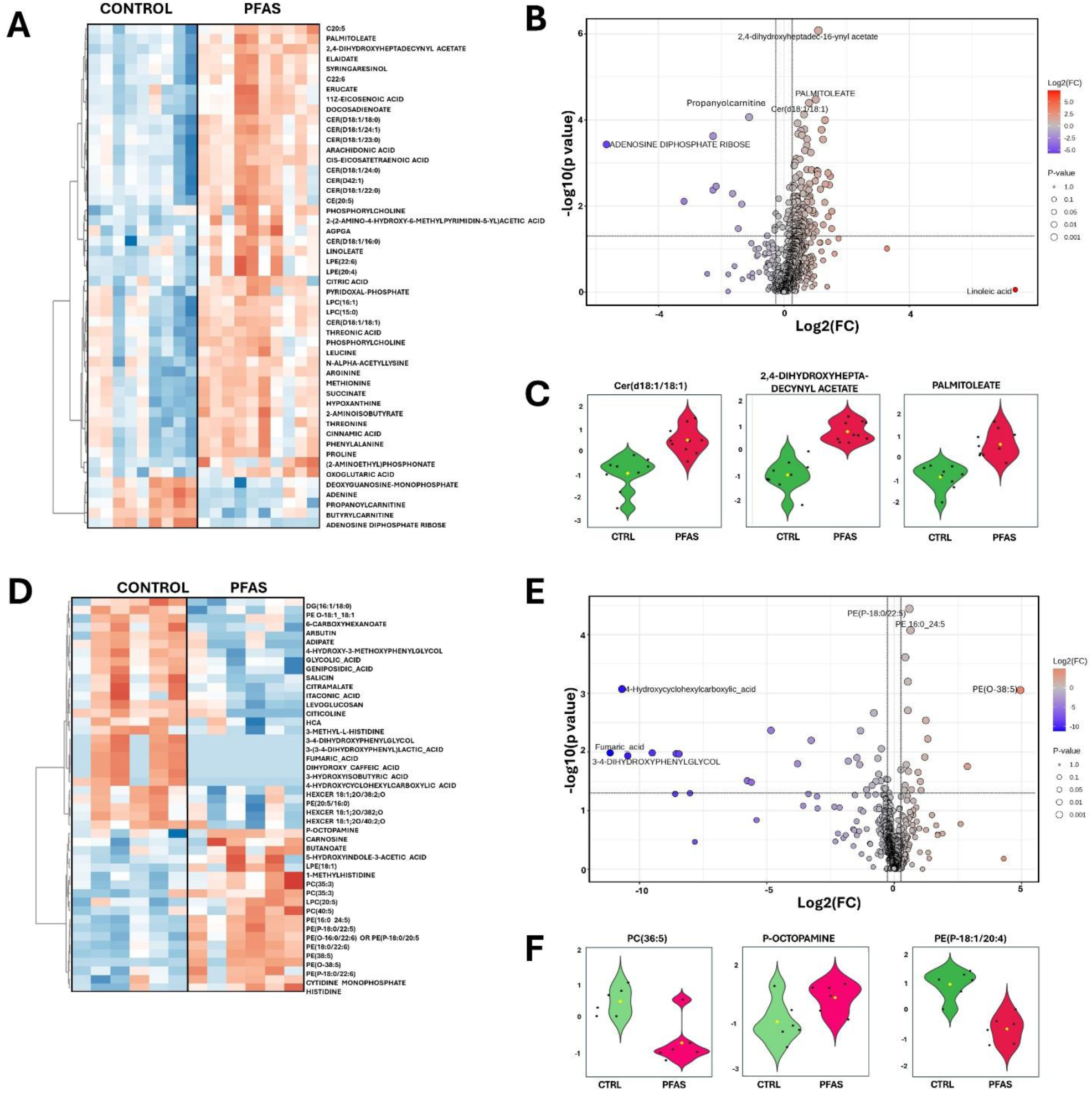
A) Heatmap showing significantly altered metabolites in female mice brain, B) Volcano plot showing differential metabolites in female mice brain (PFAS-exposed vs control). (C) Selected examples of significantly altered metabolites in female mice brain. D) Heatmap showing significantly altered metabolites in male mice brain, B) Volcano plot showing differential metabolites in male mice brain (PFAS-exposed vs control). (C) Selected examples of significantly altered metabolites in male mice brain.

**Figure 4.**
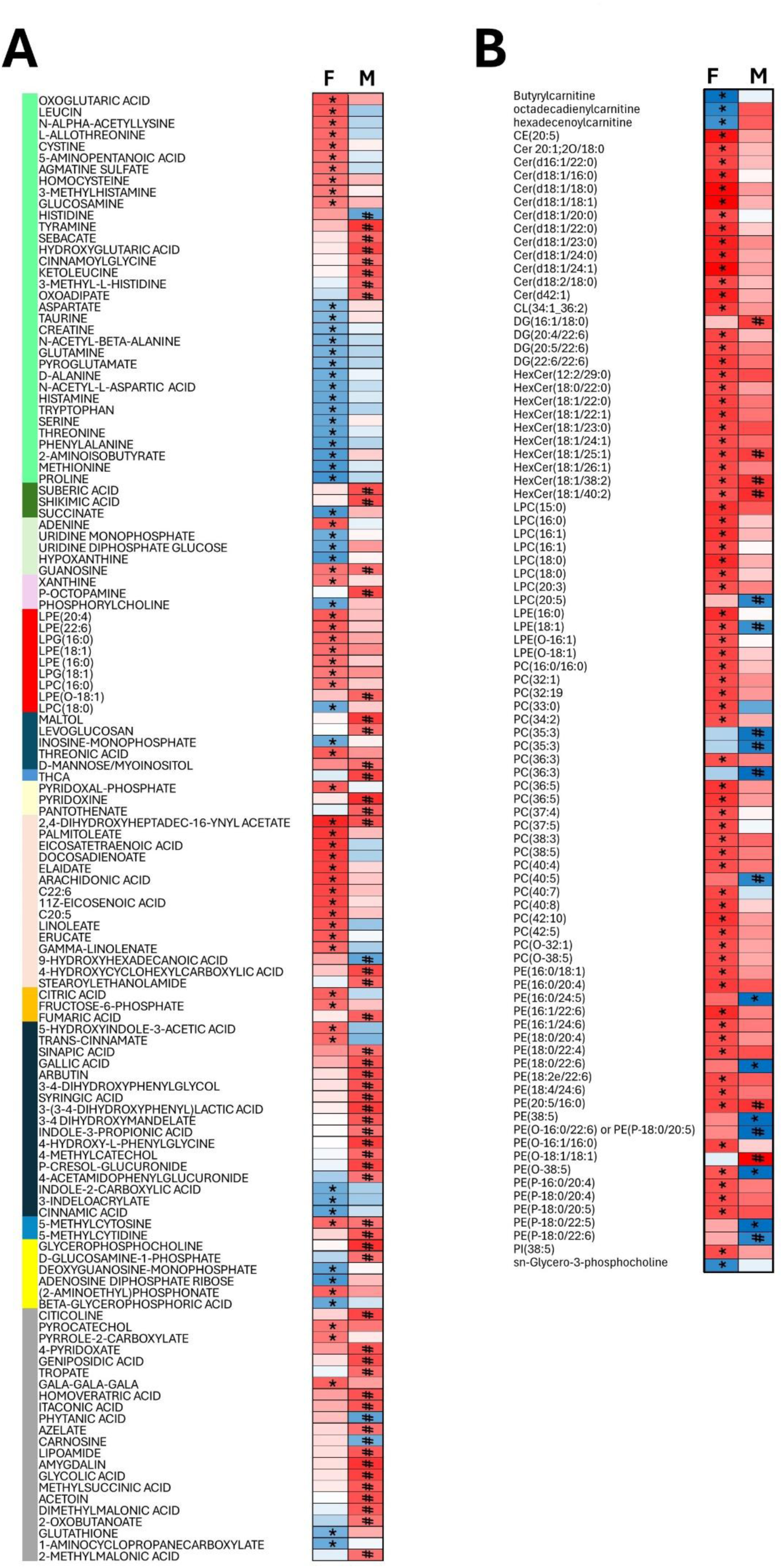
Fold changes in A) polar and semipolar metabolites B) in lipids profiles in female and male female mice * p<0.05, FDR<0.1, # p<0.05.

In contrast, male mice exhibited substantially fewer metabolic alterations in the brain. Most observed changes reached nominal statistical significance but did not survive false discovery rate (FDR) correction. Among these, five phosphatidylethanolamines were downregulated, while three metabolites—4-hydroxycyclohexylcarboxylic acid, glycolic acid, and 4-hydroxy-3- methoxyphenylglycol—were upregulated. In addition to the identified metabolites and lipids, a large number of unidentified compounds showed significant differences in both female and male mouse brains.

Next, we analyzed partial correlations among brain metabolites that showed statistically significant differences between control and PFAS-treated mice. Because only a limited number of metabolites survived FDR correction in males, this analysis was performed exclusively for female mice, separately for control and PFAS-exposed groups. As shown in Figure 5, the partial correlation patterns differed markedly between the two conditions. In control animals, metabolites formed tightly clustered networks with two main clusters: one consisting primarily of amino acids, closely associated with some lipids (mainly hexylceramides) and free fatty acids, and a second cluster composed mainly of phospholipids. In PFAS-treated mice, this organization was disrupted, resulting in three more loosely connected clusters: (1) phospholipids, (2) amino acids and carnitines, and (3) free fatty acids and other lipids, along with some nucleotides.

**Figure 5.**
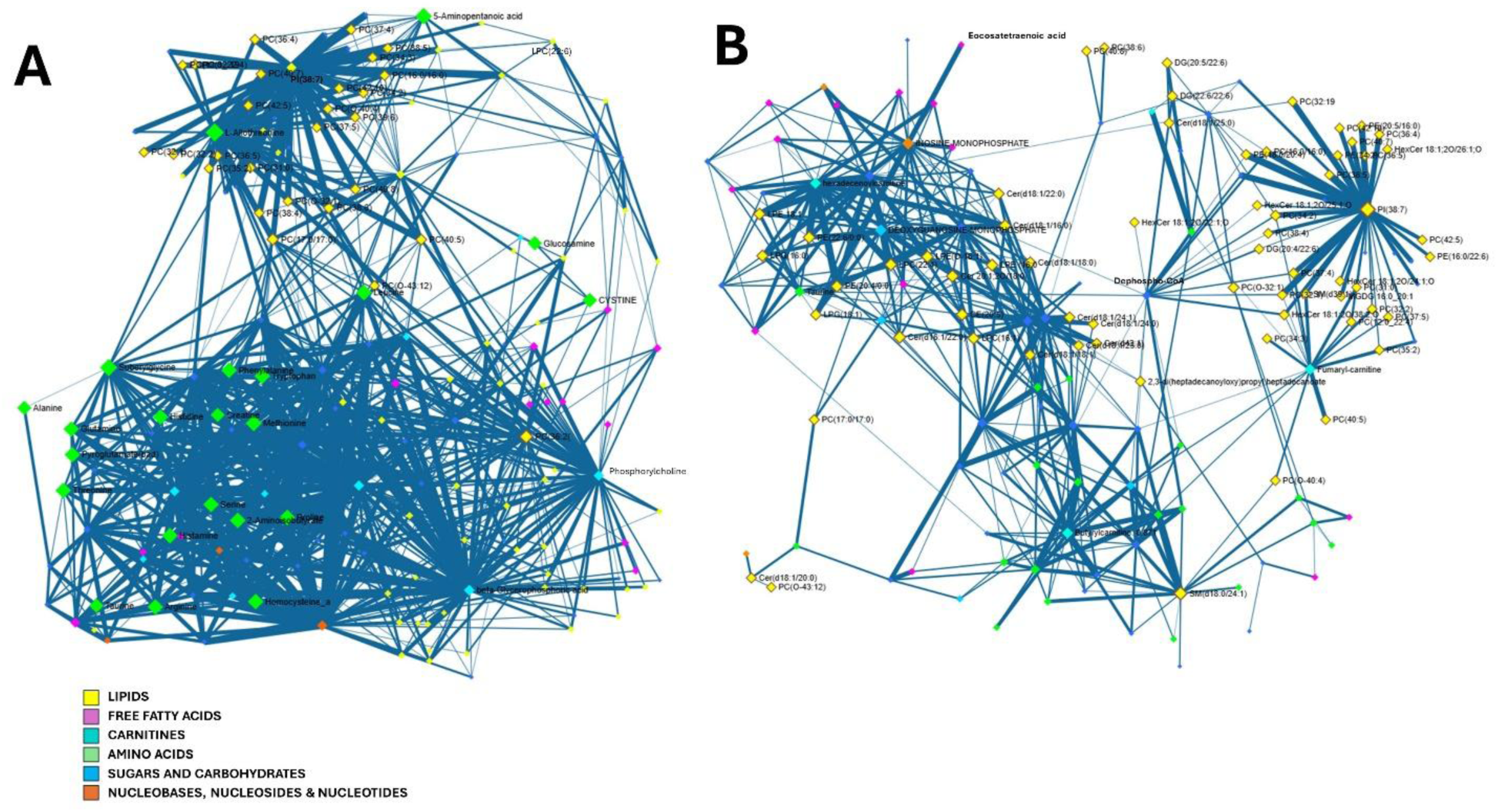
Partial correlation network of A) control female mice brain metabolites and B) PFAS treated female mice brain metabolites. Metabolites showing significant differences (p<0.05, FDR<0.1) were chosen for the plot, with p value threshold for the correlation <0.05.

### 3.3. Pathway analysis and enrichment analysis

Pathway analyses were performed separately for female and male mice and for plasma and brain samples (Figure 6, Supplementary Table 4).

**Figure 6.**
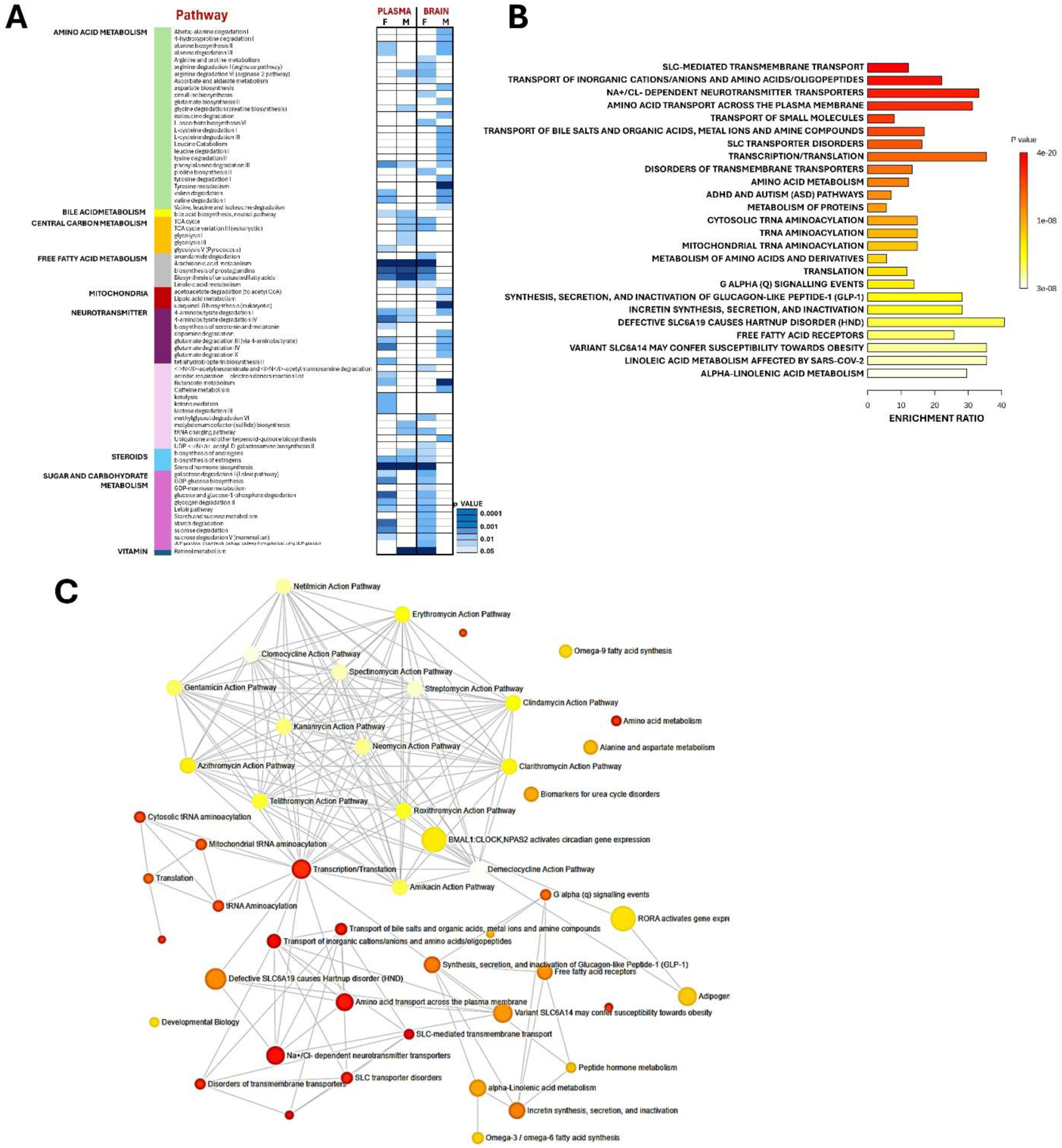
A) Significantly altered pathways based on functional analysis in female and male mice in both plasma and in the brain, B) Enrichment analysis showing top 25 altered pathways in female mice (brain), based on identified metabolites and C) network visualisation of the enriched pathways in brain of female mice.

In plasma, female mice showed a larger number of dysregulated pathways compared with male mice. Several pathways were altered in both sexes, including steroid hormone biosynthesis, bile acid biosynthesis, arachidonic acid metabolism, biosynthesis of unsaturated fatty acids, biosynthesis of prostaglandins, biosynthesis of estrogens, and 4-aminobutyrate degradation. These shared pathways mainly relate to lipid metabolism, hormone signaling, and neurotransmitter metabolism. In female mice, additional pathways associated with amino acid and carbohydrate metabolism were dysregulated, including the biosynthesis of serotonin and melatonin. In male mice, enriched pathways mainly clustered around central carbon metabolism (glycolysis and the TCA cycle), lipid metabolism, and amino acid degradation, suggesting that PFAS exposure induces broad metabolic reprogramming affecting mitochondrial energy production, lipid homeostasis, and nitrogen metabolism.

In the brain, many of the dysregulated pathways in female mice were associated with fatty acid metabolism, mitochondrial function, amino acid and nitrogen metabolism, and central carbon and energy metabolism. Several steroid hormone biosynthesis pathways were also affected. Overall, there were 13 pathways that were affected both in the plasma and in the brain in female mice, including fatty acid and steroid metabolic pathways as well as pathways related to carbohydrate metabolism.

In male mice, dysregulated pathways in the brain included those related to amino acid metabolism and neurotransmitter metabolism, such as dopamine degradation, 4- aminobutyrate degradation, tyrosine metabolism, and tyrosine degradation, as well as pathways linked to mitochondrial metabolism, including lipoic acid metabolism, ubiquinol-8 biosynthesis, and acetoacetate degradation. Only two pathways were altered both in the plasma and in the brain (4-aminobutyrate degradation, phenylalanine degradation).

Overall, several common metabolic themes emerged across plasma and brain, particularly involving lipid metabolism, steroid hormone biosynthesis, amino acid metabolism, and central energy metabolism. These shared alterations suggest that PFAS exposure may induce systemic metabolic changes affecting mitochondrial function, lipid homeostasis, and neurotransmitter-related pathways, with tissue- and sex-specific differences in the affected pathways.

Additionally, for female mice we performed enrichment analysis using the RaMP-DP pathway database based on significantly altered metabolites across all three analytical platforms (Figure 4B, Supplementary Table 5). The most affected pathways included several processes related to transmembrane transport, particularly Na⁺/Cl⁻-dependent neurotransmitter transporters, as well as amino acid metabolism and aminoacyl-tRNA biosynthesis pathways.

## 4. Discussion

Our findings demonstrate that early-life PFAS exposure is associated with broad metabolic alterations in both plasma and the brain, affecting pathways related to energy metabolism, lipid homeostasis, amino acid metabolism, and redox balance. The metabolomic changes observed in plasma indicate that PFAS exposure induces systemic metabolic reprogramming, which may influence brain metabolism either directly through PFAS accumulation in neural tissue or indirectly through alterations in circulating metabolic substrates and signaling molecules. The observed sex-specific patterns suggest differential susceptibility of male and female mice to PFAS-induced metabolic stress, providing a potential molecular framework for previously reported sex-dependent neurodevelopmental outcomes.^39^

The environmental relevance of the exposure model should be considered in the context of species-specific PFAS toxicokinetics and internal tissue concentrations. Although the PFAS concentrations administered in drinking water exceeded those typically detected in environmental drinking water, they were on the level measured in contaminated areas.^40^ In addition, higher administered doses are commonly required in mice because of their generally faster elimination of several PFAS compared with humans. In the present study, plasma PFAS concentrations exceeded those generally reported for the human general population and were more comparable to levels observed in occupationally exposed populations. Importantly, however, PFAS concentrations measured in mouse brain tissue were within a range comparable to those reported in human brain samples.^41^ These findings highlight the importance of evaluating internal tissue concentrations rather than relying solely on administered doses when assessing the human relevance of experimental PFAS exposure. The observed metabolic alterations may therefore provide insight into biological responses associated with PFAS accumulation in the brain, although differences in exposure duration, toxicokinetics, and species-specific susceptibility should be considered when extrapolating these findings to humans.

Across both plasma and the brain, several shared metabolic themes emerged, including dysregulation of lipid metabolism, steroid hormone pathways, amino acid metabolism, and central carbon metabolism. These overlapping alterations suggest that PFAS exposure may affect core metabolic networks that link peripheral metabolic regulation with brain energy homeostasis. Such systemic metabolic disturbances may influence brain development through changes in circulating lipids, amino acids, and hormone-related metabolites that are known to regulate neuronal metabolism and synaptic signaling.

Despite shared metabolic signatures, clear sex-specific differences emerged in the brain metabolome. PFAS exposure induced consistent dysregulation of amino acid metabolism in the brains of both female and male mice. Female brains showed particularly strong dysregulation of lipid metabolism and arachidonic acid (AA) pathways, along with broader alterations in carbohydrate and nucleotide metabolism. In contrast, male brains exhibited more pathway-specific alterations, primarily affecting neurotransmitter-related metabolism and mitochondrial pathways. These sex-dependent metabolic responses may reflect differences in mitochondrial vulnerability, lipid signaling, hormonal regulation, or compensatory metabolic capacity between males and females.

In plasma, the metabolic response also showed sex-specific characteristics, with male mice exhibiting a larger number of significantly altered metabolites, particularly reflecting widespread lipid downregulation and alterations in central carbon metabolism. These plasma changes may indicate systemic shifts in energy utilization and lipid transport that could influence the availability of metabolic substrates for the brain. Female plasma samples showed fewer significantly altered metabolites, although this may partly reflect the smaller number of samples available for plasma samples. Nevertheless, pathway analyses in plasma indicated dysregulation of lipid signaling, steroid hormone metabolism, and neurotransmitter-related pathways, which overlapped with several pathways affected in the brain. This overlap suggests that PFAS exposure may induce coordinated metabolic changes across tissues, potentially reflecting integrated responses within the brain–periphery metabolic axis.

In female plasma samples, strong upregulation of 3-methoxytyrosine was observed. This metabolite is linked to catecholamine metabolism and is commonly associated with dysregulation of aromatic L-amino acid decarboxylase (AADC), an enzyme involved in dopamine and serotonin biosynthesis. Elevated levels of this metabolite may therefore reflect alterations in catecholaminergic neurotransmitter pathways, suggesting that PFAS exposure may influence monoaminergic signaling through both systemic and central metabolic mechanisms. The pronounced disruption of AA metabolism observed in female brains is particularly relevant given the central role of AA in synaptic signaling, neurotransmission, and neuroprotection. These findings are consistent with the pathway analysis results in plasma, where arachidonic acid metabolism and prostaglandin biosynthesis were also significantly enriched, suggesting that PFAS exposure may influence systemic lipid signaling pathways that extend to the brain. Dysregulated AA metabolism has been linked to neuroinflammation and neurodegenerative diseases and it can influence neuronal development, synaptic plasticity, and mood disorders such as bipolar disorder.^42, 43^ Similar alterations in AA metabolism following PFAS exposure have been reported in the brains of male mice, supporting the sensitivity of this pathway to PFAS-induced neurotoxicity.^15^ The changes in carbohydrate metabolism pathways observed in our study predominantly in female brains likely reflect increased energetic demand and heightened mitochondrial stress.^44^ These shifts suggest a partial reliance on glycolytic energy production to compensate for impaired mitochondrial oxidative phosphorylation. Comparable findings have been reported in *Danio rerio*, where PFOA exposure altered galactose metabolism, starch and sucrose metabolism, and pathways involved in unsaturated fatty acid biosynthesis and BCAA metabolism.^45^ In addition to metabolic pathway disruptions, enrichment analysis in female brains revealed significant alterations in pathways related to transmembrane transport, including Na⁺/Cl⁻- dependent neurotransmitter transporters. These transporters are essential for the reuptake of neurotransmitters such as dopamine, serotonin, and GABA, and their activity is tightly coupled to membrane lipid composition, ion gradients, and mitochondrial energy availability. Disruption of these transport systems may therefore reflect functional consequences of PFAS-induced membrane remodeling and bioenergetic stress, potentially impairing synaptic neurotransmitter clearance and signaling fidelity. Alterations in Na⁺/Cl⁻-dependent transporter pathways are also consistent with the observed lipidomic changes, as membrane phospholipid composition and fluidity directly regulate transporter conformation and activity. Together, these findings suggest that PFAS exposure in female brains not only affects metabolic and lipid homeostasis but may also compromise neurotransmitter transport processes critical for synaptic function and neuronal communication.

PFAS exposure also induced pronounced lipidomic alterations in the brains of female mice, consistent with enhanced membrane turnover and compensatory remodeling, and potentially linked to altered myelination. Specifically, reduced glycerophosphorylcholine (GPC) together with increased phospholipids, including lysophosphatidylcholines (LPCs), phosphatidylcholines (PCs), and phosphatidylethanolamines (PEs), suggests increased membrane remodeling and reduced phospholipid catabolite availability. Choline also showed a trend toward reduction, although this change did not reach statistical significance. The upregulation of LPCs, together with increased free fatty acids, particularly arachidonic acid (AA) and docosahexaenoic acid (DHA), is indicative of phospholipase A2–driven membrane phospholipid cleavage, likely triggered by oxidative and/or inflammatory stress.^46^ Concurrent increases in PC and PE suggest compensatory membrane repair or rebuilding in response to this lipid damage, consistent with the observed reduction in GPC. These lipid alterations may be mechanistically linked to disrupted myelination or remyelination processes during early neurodevelopment. The myelin sheath has a highly specialized lipid composition, and specific lipid classes have been associated with both de- and remyelination. ^47^ Increased polyunsaturated glycerophospholipids have been linked to demyelination in the corpus callosum, whereas elevated free fatty acids may reflect active remyelination processes. ^47^ In line with this interpretation, epidemiological studies have shown that maternal PFAS exposure is associated with alterations in children’s white matter microstructure in a sex-specific manner.^48^ While PFHxS and PFOS isomers have been associated with corpus callosum alterations predominantly in males, PFHxS has been linked to changes in the superior and inferior longitudinal fasciculi, and PFOS isomers to alterations in pyramidal fibers and the inferior longitudinal fasciculus in females. These findings suggest that prenatal PFAS exposure may interfere with white matter development, potentially through disrupted myelin formation or regeneration.^48^ Supporting this hypothesis, animal studies have demonstrated that PFAS exposure severely impairs remyelination.^28, 49^ Notably, abnormalities in corpus callosum white matter are a prominent feature of neurodevelopmental disorders such as attention-deficit/hyperactivity disorder (ADHD) and are thought to contribute to associated cognitive and motor impairments. .^48^.^48, 50^ Together, these observations suggest that PFAS-induced lipid dysregulation— particularly in the female brain—may represent a biologically plausible pathway linking prenatal exposure to altered white matter development and subsequent neurodevelopmental outcomes.

Additionally, reduced acetylcarnitine levels in female brain indicate mitochondrial stress and impaired acetyl-CoA handling, a finding further supported by increased ceramide levels, which are known mediators of mitochondrial dysfunction and inflammatory signaling. In parallel, downregulation of purine and pyrimidine nucleotides, together with reduced ADP-ribose derivatives, suggests impaired nucleotide synthesis and diminished NAD⁺-dependent metabolic capacity. This together with the concomitant depletion of glutathione and N-acetyl-aspartate points to compromised antioxidant defence and neuronal mitochondrial dysfunction. Collectively, these metabolic changes support a model in which PFAS exposure induces chronic bioenergetic stress and oxidative damage in the brain, driving imbalanced membrane remodeling and reduced neuronal metabolic resilience. These brain metabolic alterations align with plasma pathway changes involving central carbon metabolism and amino acid degradation, further supporting the hypothesis that PFAS exposure disrupts mitochondrial bioenergetics at both systemic and neural levels.

In contrast, metabolic changes in the male brain were generally less extensive but more pathway-specific, particularly affecting neurotransmitter-related metabolism. These findings parallel the plasma pathway results showing enrichment of amino acid degradation and central carbon metabolism, suggesting coordinated metabolic adjustments affecting neurotransmitter precursor availability. Also, while the changes in brain lipidome were not as strong as observed in the female brains, they trend of change was similar. Notably, dopamine degradation and tyrosine metabolism were strongly impacted. Tyrosine serves as a precursor for catecholamine neurotransmitters, including dopamine, norepinephrine, and epinephrine. Dopamine plays a critical role in motor control, reward processing, and cognitive function, and its degradation is mediated primarily by monoamine oxidase and catechol-O-methyltransferase, as well as by dopamine transporter mediated reuptake. These processes are sensitive to oxidative stress, and dysregulation has been linked to neurodegenerative disorders such as Parkinson’s disease.^51–53^ Experimental studies have shown that PFAS, particularly PFOS, can reduce dopamine levels in the brain.^54^ In line with these findings, alterations in tyrosine metabolism and ubiquinol-8 biosynthesis observed here further suggest impaired antioxidant and mitochondrial function in male brains. Additionally, butanoate metabolism was disrupted in males. Butanoate functions as both an energy substrate and a signaling molecule in the brain, influences the gut–brain axis, inhibits histone deacetylases to regulate gene expression, and supports neuroprotection and cognitive function. ^41^ Disruption of this pathway may therefore contribute to PFAS-induced neurofunctional impairments affecting neurotransmitter precursor availability.

Partial correlation analysis revealed a marked reorganization of metabolite–metabolite relationships in PFAS-exposed female mice, suggesting disruption of coordinated metabolic regulation rather than isolated changes in individual metabolites. In control animals, metabolites formed tightly connected clusters, with amino acids closely integrated with specific lipid species and a distinct phospholipid cluster, consistent with normal coupling between amino acid metabolism, lipid homeostasis, and membrane turnover. In contrast, PFAS exposure fragmented this organization into more weakly connected clusters, with separation of phospholipids, amino acids/carnitines, and free fatty acids. The clustering of carnitines with amino acids may reflect altered mitochondrial fatty acid transport and energy metabolism, while the dissociation of free fatty acids from phospholipids suggests impaired lipid remodeling.

Together, the integrated plasma and brain metabolomics results indicate that PFAS exposure induces multi-organ metabolic perturbations, suggesting that both direct neurotoxic effects and systemic metabolic disturbances may contribute to PFAS-associated neurodevelopmental outcomes.

A key strength of this study is the integrated metabolomic and lipidomic profiling of the brain, which enabled identification of coordinated PFAS-induced disruptions in energy metabolism, lipid homeostasis, redox balance, and neurotransmitter-related pathways. The sex-stratified analysis revealed distinct metabolic vulnerability patterns in female and male brains, highlighting pronounced lipid remodeling and arachidonic acid dysregulation in females and more pathway-specific effects on neurotransmitter and mitochondrial metabolism in males. These sex-dependent molecular signatures provide mechanistic insight into previously reported sex-specific neurodevelopmental outcomes following PFAS exposure. The prenatal exposure model and the consistency of affected pathways with prior experimental and epidemiological studies further support the biological relevance of the findings. Several limitations should be acknowledged. The relatively small sample size reduces statistical power and may limit detection of more subtle effects. In addition, the use of NOD mice, which have strain-specific metabolic and immunological characteristics, may influence susceptibility to PFAS exposure and limits generalizability to other strains or humans. The study is also primarily correlative, as functional assessments of mitochondrial activity, neurotransmitter signaling, or white matter integrity were not performed. Finally, analyses were conducted in whole-brain tissue, precluding region- or cell-type-specific interpretation.

### Summary

In summary, prenatal PFAS exposure induces broad and sex-dependent metabolic reprogramming in the developing brain, affecting pathways linked to mitochondrial function, lipid remodeling, oxidative stress, and neurotransmitter metabolism. The pronounced lipidomic and bioenergetic disturbances observed in female brains, alongside more targeted neurotransmitter-related changes in males, support a model in which PFAS exposure compromises neuronal metabolic resilience in a sex-specific manner. Our findings also indicate that prenatal and early-life PFAS exposure induces persistent network-level metabolic alterations, highlighting disrupted metabolic integration as a potential mechanism underlying PFAS-associated developmental effect. Despite limitations related to sample size, strain specificity, and lack of functional validation, these findings provide a mechanistic framework linking prenatal PFAS exposure to altered brain development and highlight the importance of considering sex as a critical biological variable in PFAS neurotoxicity research.

## Acknowledgements

This study was supported by the Swedish Research Council (grant no. 2020-03674 to T.H and M.O.), by the Swedish Knowledge Foundation (grant no. 20220122, to T.H. and M.O.).and by the Inflammation in human early life: targeting impacts on life-course health (INITIALISE) consortium funded by the Horizon Europe Program of the European Union under grant agreement 101094099 to M.O. and by the Swedish Knowledge Foundation (grant no. 20220122, to T.H. and M.O.).

Views and opinions expressed are those of the author(s) only and do not necessarily reflect those of the European Union. Neither the European Union nor the granting authority can be held responsible for them.

